# Genetic Variation Regulates Opioid-Induced Respiratory Depression in Mice

**DOI:** 10.1101/2020.06.15.152249

**Authors:** Jason A. Bubier, Hao He, Vivek M. Philip, Tyler Roy, Christian Monroy Hernandez, Kevin D. Donohue, Bruce F. O’Hara, Elissa J. Chesler

## Abstract

In the U.S., opioid prescription for treatment of pain nearly quadrupled from 1999 to 2014, leading to an epidemic in addiction and overdose deaths. The most common cause of opioid overdose and death is opioid-induced respiratory depression (OIRD), a life-threatening depression in respiratory rate thought to be caused by stimulation of opioid receptors in the inspiratory-generating regions of the brain. Studies in mice have revealed that variation in opiate lethality is associated with strain differences, suggesting that sensitivity to OIRD is genetically determined. We first tested the hypothesis that genetic variation in inbred strains of mice influences the innate variability in opioid-induced responses in respiratory depression, recovery time and survival time. Using the founders of the advanced, high-diversity mouse populations, the Collaborative Cross (CC) and Diversity Outbred (DO), we found substantial sex and genetic effects on respiratory sensitivity and opiate lethality. To define genetic modifiers of OIRD, we then used the high precision DO population treated with morphine to map and identify quantitative trait loci (QTL) for respiratory depression, recovery time and survival time. Trait mapping and integrative functional genomic analysis in GeneWeaver has allowed us to implicate *Galnt11*, an N-acetylgalactosaminyltransferase, as a candidate gene that regulates OIRD.

## Introduction

One in every four individuals treated for pain with prescription opioids such as morphine becomes addicted and progresses to illicit synthetic opioid use^1^. The non-uniform composition, dosing and administration of more potent synthetic street opioids including fentanyl frequently leads to overdose. Remedial measures such as treatment with naloxone, a competitive antagonist of the opioid receptor mu 1 (OPRM1), the principal target of opioids, are often unsuccessful due to the higher potency of synthetic opioids^2-4^. Therefore, novel approaches toward understanding opioid addiction, overdose and remediation are essential.

The most frequent cause of overdose death due to opioids is opioid-induced respiratory depression (OIRD). Opioids such as morphine depress the hypoxic ventilatory response in the brainstem by affecting the chemosensitive cells that respond to changes in the partial pressures of carbon dioxide and oxygen in the blood^5^. Understanding the underlying molecular mechanisms that control the respiratory responses to opioids may provide insight into alternative therapies for opioid overdose by treating the respiratory depression directly, independent of, or together with opioid receptor antagonism. Variable responses in OIRD have been well documented in both humans and mice^6^, and likely occur through diversity in the molecular pathways within the ventilatory processing centers of the brainstem. Our goal is to utilize this genetic diversity to define genetic modifiers of OIRD in mice.

Genetic diversity in laboratory mice is a powerful tool for dissecting the genetic and molecular components of complex traits such as the response to opioids. In mice, genetic variation has been identified in the opioid receptors^7-11^, in genes and pathways associated with the anti-nociceptive effects of morphine^12-17^ and morphine withdrawal^18^, and in other behavioral responses to opioids^16,19,20^. Other studies have characterized the median lethal dose (LD_50_) of morphine across mouse strains^21-23^. One study found that a strain harboring an OPRM1 hypomorph (CXB7), in which OPRM1 gene function is reduced, have a much higher LD_50_ for morphine than strains with intact mu-opioid^8,24^, confirming the importance of OPRM1 in opioid-induced lethality and revealing strain diversity in the mu-opioid receptor locus. Together, these studies show that variation in response to opioids is associated with differences in strain, indicating that responses to opioids are genetically influenced phenotypes. However, these studies were generally conducted on only a few strains of male mice with survival as the only endpoint, and thus did not characterize known sex differences in opioid sensitivity^25^ and were not designed to define the genetic loci or the underlying biological mechanisms responsible for the strain variation in complex opioid responses, such as respiratory depression.

Respiratory phenotyping in mice generally includes measures of ventilation mechanics such as respiratory frequency and tidal volume using conventional plethysmography. However, plethysmography is labor intensive and the duration over which the mouse can be monitored is limited because the animal is confined and its movements are restricted. Further, monitoring the rapid response to drugs by plethysmography is invasive because it requires the surgical implantation of a cannula, catheter or other port into the mouse in order to administer the compounds. Here, we have taken the novel approach using signals obtained from a piezo electric sleep monitoring system^26^ to measure respiratory phenotypes. This system offers a non-invasive, high-throughput technology to study respiratory depression in response to opioids in mice in a home-cage setting. The piezo technology was adapted to estimate average respiratory rates over specified time intervals for characterizing patterns associated with respiratory depression. This allows for the determination of respiratory depression and time to recovery or cessation of respiration. This system was previously validated and used by us in genetic studies of sleep in the Collaborative Cross (CC)^27,28^, BXD ^18^, and other mouse populations^29^.

In this study, we investigated the effect of genetic variation on OIRD and lethality in the founders of the advanced, high-diversity mouse populations, specifically the CC and Diversity Outbred (DO) populations. After determining that the quantitative respiratory phenotypes were heritable and that genetic mapping was therefore feasible, we mapped key respiratory traits using a population of 300 DO mice of both sexes and our advanced piezo electric sleep monitoring system^26^. This enabled us to reveal a previously undiscovered potential mechanism of variability in OIRD, which was further evaluated through analysis of sequence variation, gene expression and conservation of protein domains.

## Results

### Development of a high-throughput method for measuring respiratory depression in response to opioids

The PiezoSleep system was adapted for use as an in depth, high-throughput, continuous monitoring approach for key respiratory responses to morphine. The piezoelectric transducer in the cage floor produces voltage in direct proportion to the pressure applied to the cage floor. The pressure signal is amplified, filtered, and digitized for processing to extract signal features related to irregular wake-like activity and regular sleep-like motions primarily resulting from respiration (thorax pressure directly on the cage floor or pressure variations being transferred through the legs). Although the original PiezoSleep system only provides respiratory estimates during sleep, the algorithm was adapted to find periods of low activity and estimate respiratory rates when breathing signals were detected. These were averaged over larger intervals to provide an average respiration rate every 12 minutes to obtain a baseline rate (24 hours pretreatment) that could be compared to post-treatment responses. This enabled a quantitative evaluation of respiratory depression, survival time and recovery time (Fig. 1).

**Figure 1.**
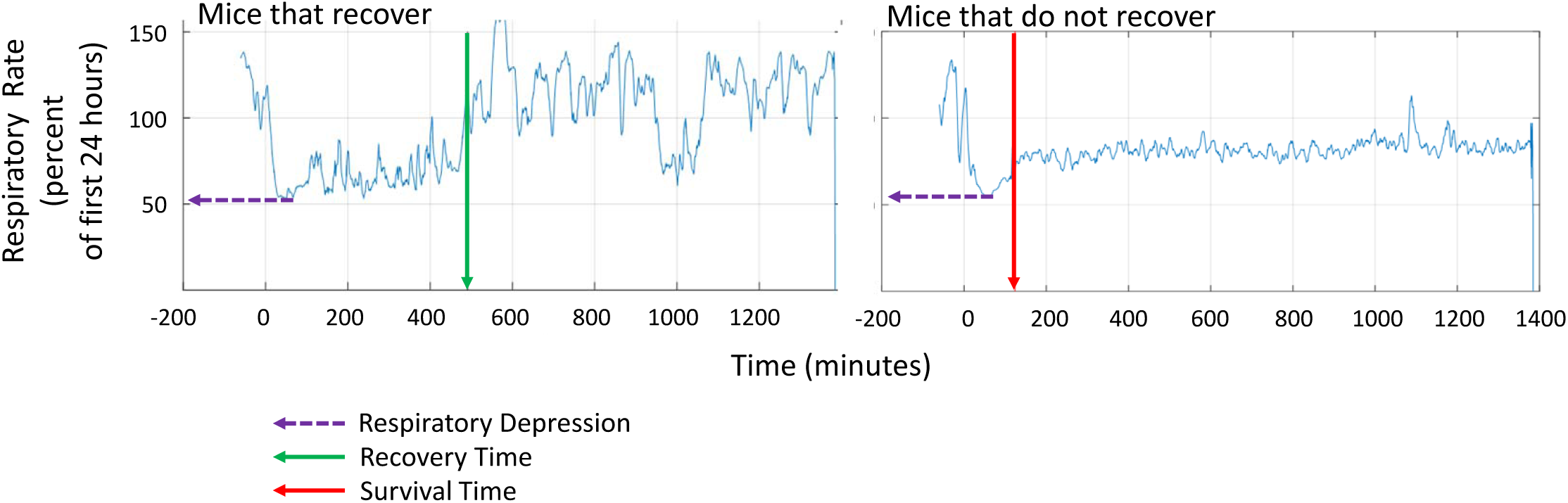
PiezoSleep output for a mouse that recovers and a mouse that fails to recover after opioid treatment. Baseline respiratory rate is first established for 24 hr. Time 0, is the time at which morphine is administered. The *blue line* represents the derived respiratory rate trajectories relative to the baseline (defined as the average respiratory rate over the first 24 hours (set at 100%)). Respiratory depression is defined as the lowest percentage of baseline reached after morphine treatment (*purple dotted arrow*). For a mouse that recovers, the *green vertical arrow* indicates recovery time when the respiratory rate returns to baseline. For a mouse that does not recover, the *red vertical arrow* indicates survival time when breathing stops (i.e., piezo output becomes machine noise never returning to baseline).

### Respiratory depression, recovery time and survival time are heritable traits

A bracketing approach was used to construct a dose-response curve with a minimal number of mice. Eight inbred strains of mice (CC and DO founders) of both sexes were tested with an initial probe dose of 436 mg/kg of morphine. We found a wide range of outcomes for respiratory depression (49% to 77%), recovery times (0.05 hours to 8.61 hours) and survival times that segregate largely by strain, but also by sex (Fig. 2). Respiratory depression at the 436 mg/kg dose showed a significant effect of strain (F_strain (7,80)_ = 10.7357, p<0.0001), but no sex effect or sex × strain interaction. Using time as the independent variable, we fit a linear model that revealed significant effects of strain, sex and dose on both recovery time (F_(25,152)_ = 4.2774; p<0.0001) and survival time (F_(24,162)_ = 10.1922; p<0.0001). From these quantitative measurements, we calculated the strain intra-class correlation (ICC) as an estimate of heritability and found that respiratory depression ICC = 0.440, recovery time ICC = 0.345 and survival time ICC = 0.338. These values indicate that the traits are heritable and also amendable to genetic mapping. All strain data have been deposited in the Mouse Phenome Database (RRID:SCR_003212 ProjectID:Bubier3).

**Figure 2.**
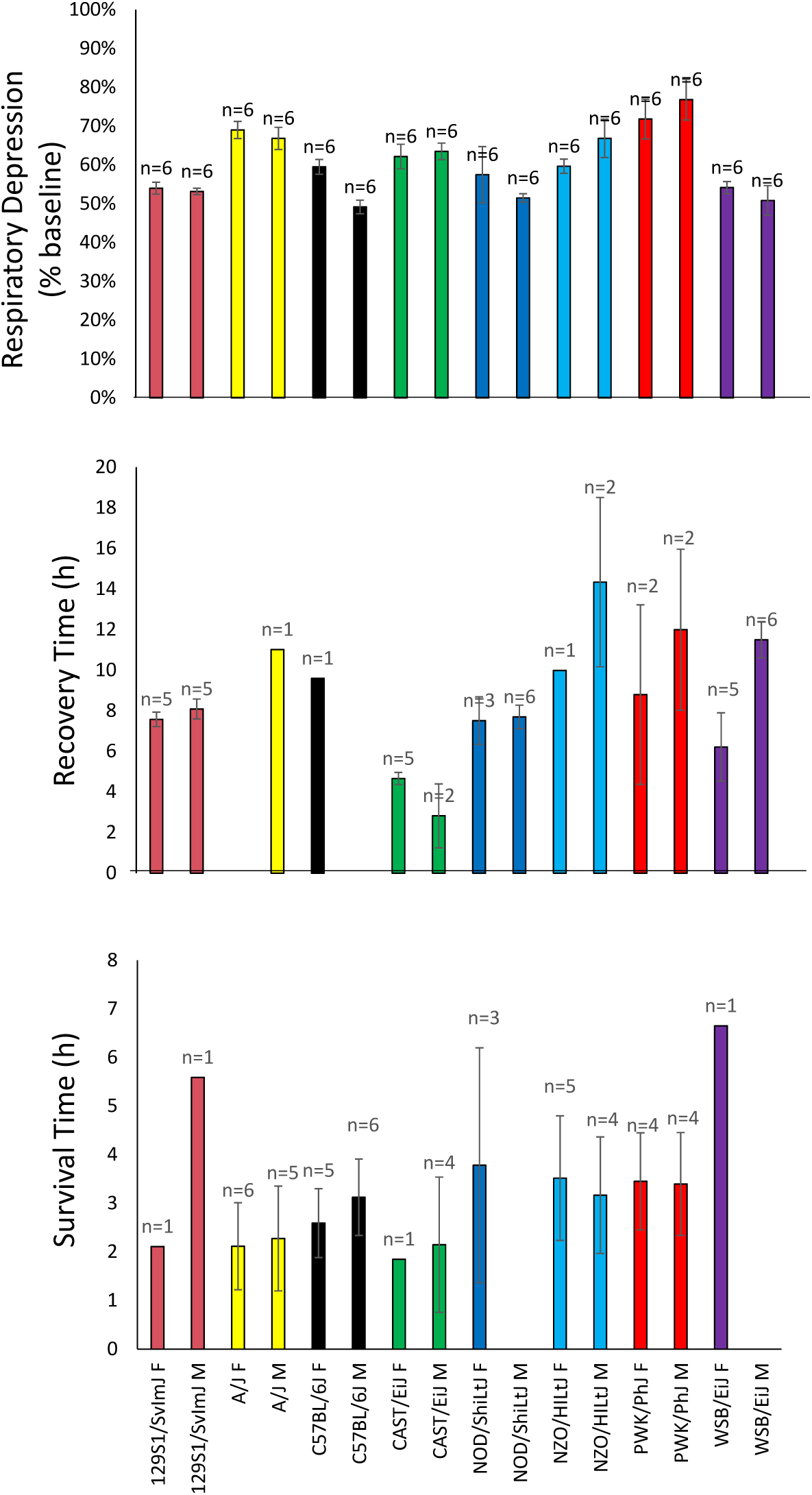
Strain- and sex-specific effect of morphine on respiratory sensitivity. Respiratory depression (top panel), recovery time (middle panel) and survival time (bottom panel) were determined using the probe dose of 436 mg/kg and the PiezoSleep system. The traits of recovery time or survival time are censored such that a mouse does not appear in both graphs as each mouse displays only one of these two phenotypes. Empty bars indicate that no mice fell into this category (i.e., either all recovered or none recovered).

### LD_50_ determination

Published data reveal a variety of morphine LD_50_ values for mice. In prior studies, using C57BL/6BY and CXB7, the LD_50_ values were 436 mg/kg and 977 mg/kg, respectively^24^. Yoburn et. al have shown that the LD_50_ for different populations of outbred Swiss-Webster range from 313 mg/kg to 745 mg/kg.^23^ Based upon this literature, we selected a probe dose of 436 mg/kg. To generate the LD_50_ curve using the minimum number of mice, additional higher or lower doses of morphine were included based on the response to the probe dose until an LD_50_ could be estimated accurately, whereby doses on each side of the 50% mark were tested. Using this approach, survival curves by strain and sex were established (Fig. 3A) and the LD_50_ for each was determined (Fig. 3B). The morphine LD_50_ ranged from 212.2 mg/kg in A/J females to 882.2 mg/kg in CAST/EiJ females. There was not a consistent sex bias up or down across all strains; instead, some LD_50_ values were higher in females than males (e.g., CAST/EiJ and NOD/ShiLtJ), but higher in males than females (e.g., PWK/PhJ and WSB/EiJ).

**Figure 3.**
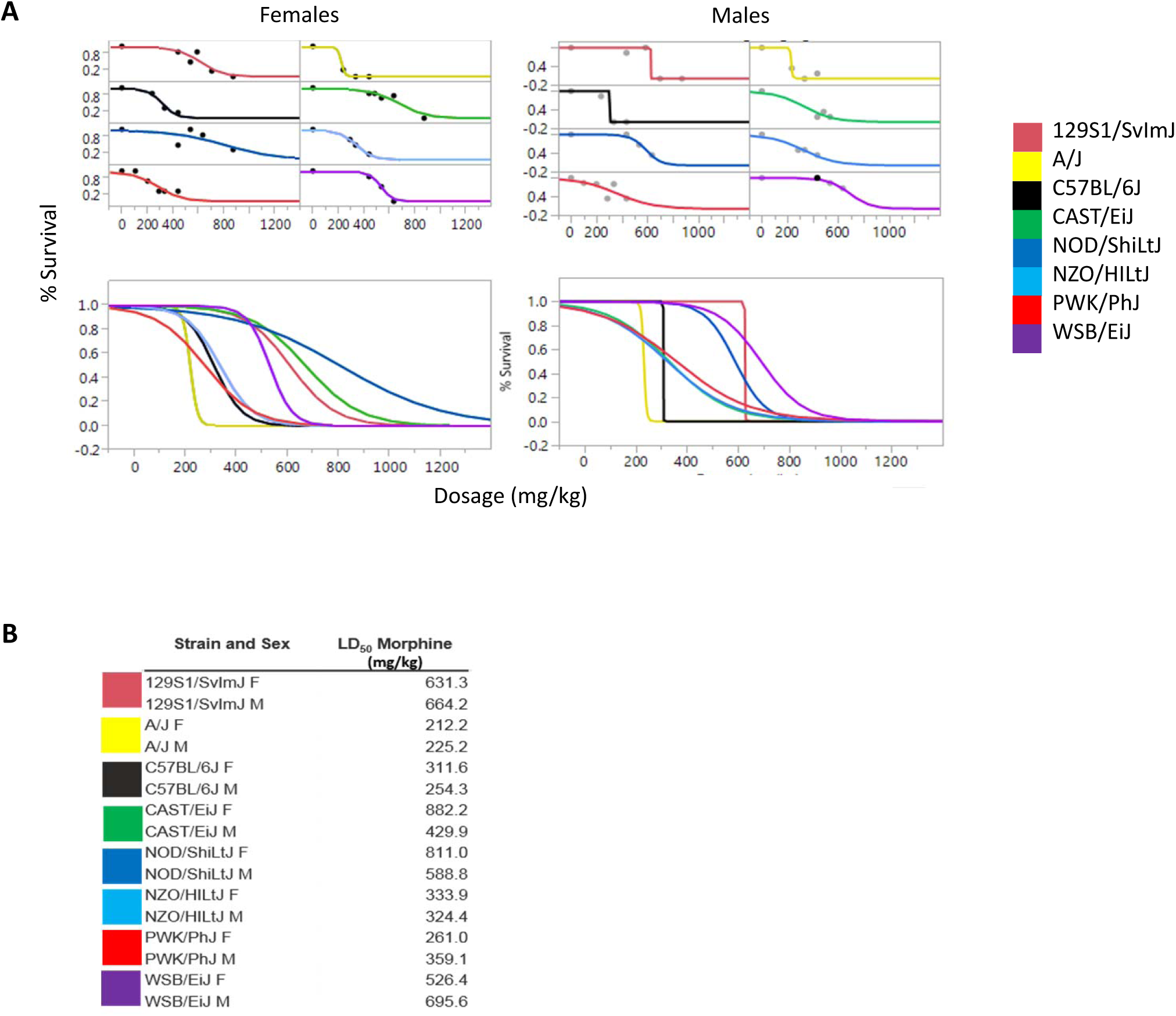
The morphine LD_50_ by strain and sex. The morphine LD_50_ was determined for each of the eight founder strains and sex using at least six mice in each group and at least three doses of morphine, with at least two doses flanking the LD_50_. (**A**) Logistic 2-paramater survival curves separated by strain and sex are shown at the top and composites of all strains separated by sex are shown at the bottom. (**B**) The LD_50_ as calculated using the drc package in R and ranged from 212 mg/kg -882 mg/kg.

### Genetic mapping in DO mice

To find genetic loci that influence OIRD, QTL mapping was performed on the respiratory phenotypes using the high diversity, high precision, DO mouse population. A probe dose of 486 mg/kg, chosen based upon the average LD_50_ of the eight founder strains and two sexes, was given to 300 DO mice, 150 of each sex. Of the 300 DO mice entered into the study, 193 (83 females, 110 males) recovered and 107 (67 females, 40 males) did not recover. The quantitative metrics for respiratory response to morphine, including respiratory depression, recovery time and survival time, all show a continuum of phenotypic diversity (Fig. 4).^30^

**Figure 4.**
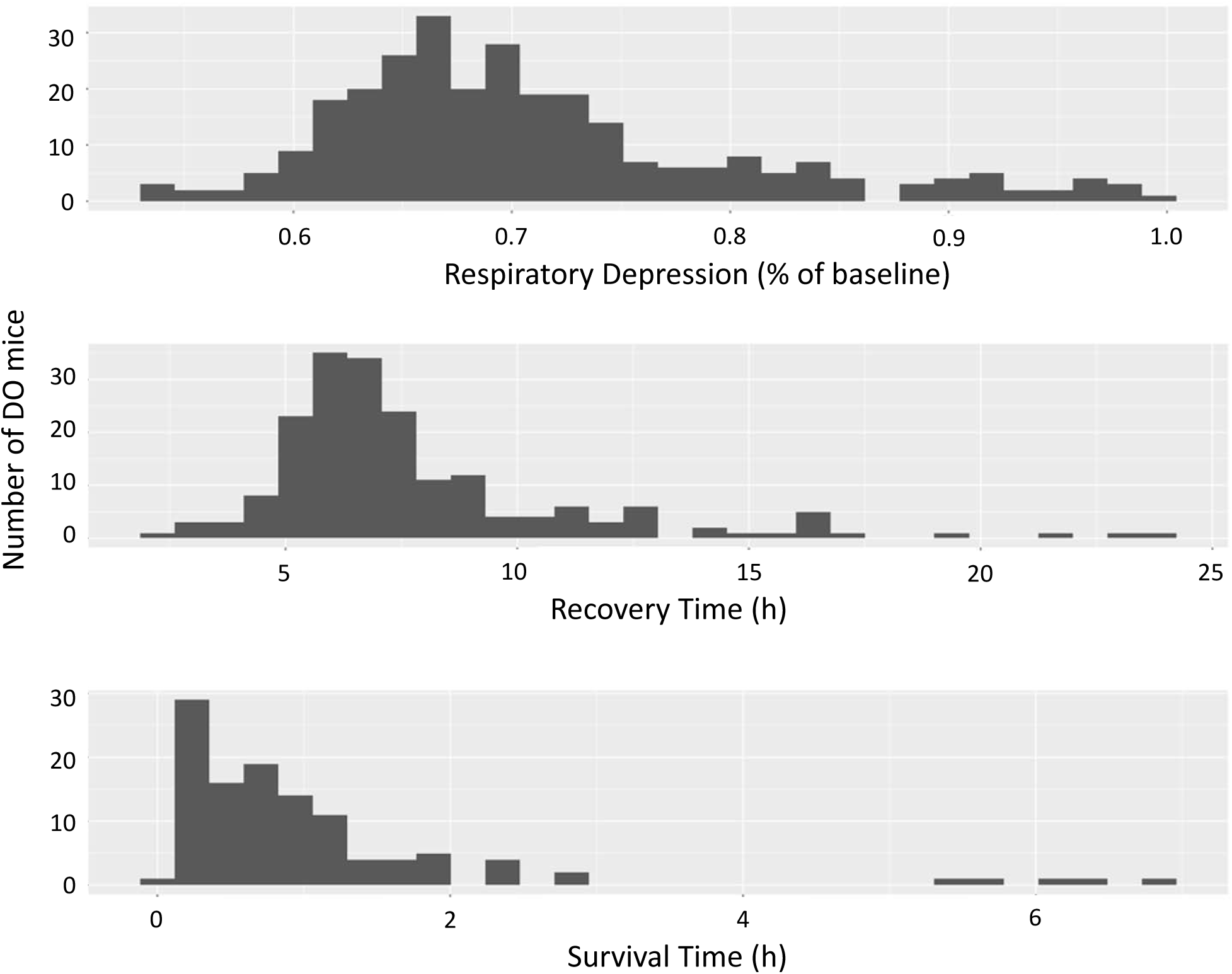
Respiratory response to morphine in Diversity Outbred mice. DO mice were given 486 mg/kg dose of morphine and the respiratory responses were determined by the Piezo Sleep system. The distribution of respiratory responses is shown for respiratory depression (top panel), recovery time (middle panel) and survival time (bottom panel).

### Respiratory response QTL

We next mapped QTL in the DO for the three respiratory traits using R/qtl2^30^ which implements an eight-state additive haplotype model. A significant QTL was identified for respiratory depression, but no genome-wide significant QTLs were detected for recovery time or survival time using these sample sizes. One suggestive QTL was identified for overall morphine survival using a COXPH model as has previously been used for QTL mapping^31^ (Fig. S1).

For the respiratory depression trait, we identified a LOD 9.2 QTL with a 1.5 LOD drop interval on Chr 5:24.30-26.25 (Fig. 5A). The 95% LOD score threshold was 7.65 for p < 0.05. The QTL is called *Rdro1* (respiratory depression, response to opioids 1). The *Rdro1* QTL is driven by strong NOD vs. WSB/EiJ allele effects (Fig. 5B). A two-state SNP association analysis identifies potential causative SNPs and results in a reduction of the 1.5 LOD drop credible interval from the peak marker (24.98-26.16 MBp), purple SNPs in Fig 5C. An ANOVA shows that the peak marker for the respiratory depression phenotype accounts for 42.8% of heritable variation in the DO population.

**Figure 5.**
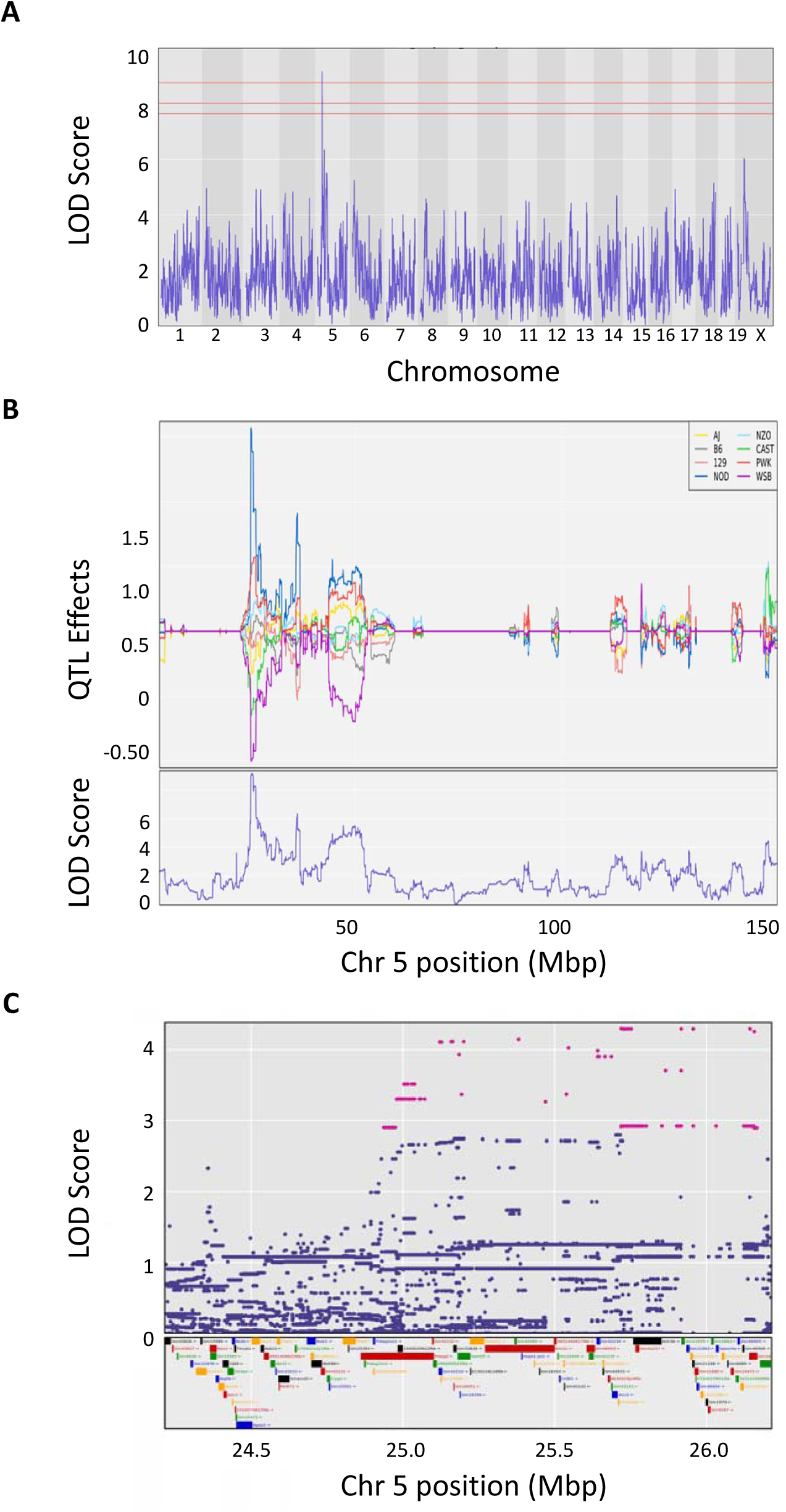
QTL mapping of respiratory depression in DO mice. (**A**) Genome wide scan for QTL regulating the phenotype of respiratory depression in 300 DO mice. (**B**) Allele effect plot of the LOD 9.2 QTL on chromosome 5 for respiratory depression showing a strong narrow peak with LOD confidence interval of 24.98-26.16 MBp. (**C**) A 2MBp interval around the peak locus on chromosome 5 showing the SNPs driving the QTL as well as the genomic features within the interval on chromosome 5.

### Identification of candidate genes

Genetic mapping studies are used to identify regions of interest containing variants that influence complex traits. To identify the relevant genes involved in complex trait regulatory mechanisms, there must be evidence of genetic polymorphisms segregating in the population that either influence protein structure or gene expression and evidence of a biological mechanism of action connecting them to the trait, such as expression in a trait-relevant tissue. We identified a 24.98-26.16 MBp credible interval containing 10,782 SNPs (including insertions and deletions) across the DO mice. More specifically using the SNP association mapping model, we found that 1,885 of these SNPs alleles were in the interval (Fig. 5B, Table S1). Of these 1,885 SNPs, eight were in coding regions of genes, 11 were in 5’UTRs, 16 were in 3’UTRs, 861 were intronic, 932 were intergenic, and 107 were in non-coding transcripts (some SNPs have multiple functions, Table S1). Non-coding variants typically influence phenotype through effects on regulation of gene expression via SNPs in regulatory regions often located in the 5’ UTR, 3’ UTR, introns or within intergenic regulator features. These potential regulatory SNPs are all located in *Speer4a, Actr3b, Xrcc2, Kmt2c, Galnt5*, and *Prkag2*. While these SNPs remain candidates for regulation of the respiratory depression phenotype, we focused on coding SNPs because their impact is more readily predictable. The eight coding SNPs lie in four genes (*Galtn11, Kmt2c, Speer4a*, and *Galnt5*). None of the coding SNPs are the type with the most deleterious effects, such as a stop loss, stop gain or coding region insertion (frameshift). All four of these protein-coding genes are expressed in the mouse pre-Bötzinger complex, a group of brainstem interneurons that control regulation of respiratory rythmogenicity^32^, and are thus mechanistically plausible. Three of these four pre-Bötzinger complex-expressed protein-coding genes contain polymorphisms that cause non-synonymous (Cn) amino acid changes. Two of the three changes (serine to threonine at amino acid 702 in *Kmt2c* and asparagine to aspartic acid at amino acid 29 in *Speer4a*) occurred in residues that are not conserved across species. The remaining gene expressed in the pre-Bötzinger complex and containing a SNP in a conserved coding region is polypeptide N-acetylgalactosaminyltransferase 11 (*Galnt11*).

### Galnt11 SNP analysis

The *Galnt11* SNP, rs37913166 C/T, changes polar serine residue 545 to a hydrophobic leucine residue (Fig. 6). This amino acid is located within a functional domain that is conserved in vertebrates (Fig. 7A), suggesting that this base change regulates a key functionality. Indeed, this change places the hydrophobic residue, which are generally buried internally, onto the surface of the protein. The 3D protein structure analysis (Fig. 7B) indicates that the non-synonymous serine-to-leucine SNP occurs in a conserved residue within the functional Ricin B lectin domain, a domain generally involved in sugar binding required for glycosylation.

**Figure 6.**
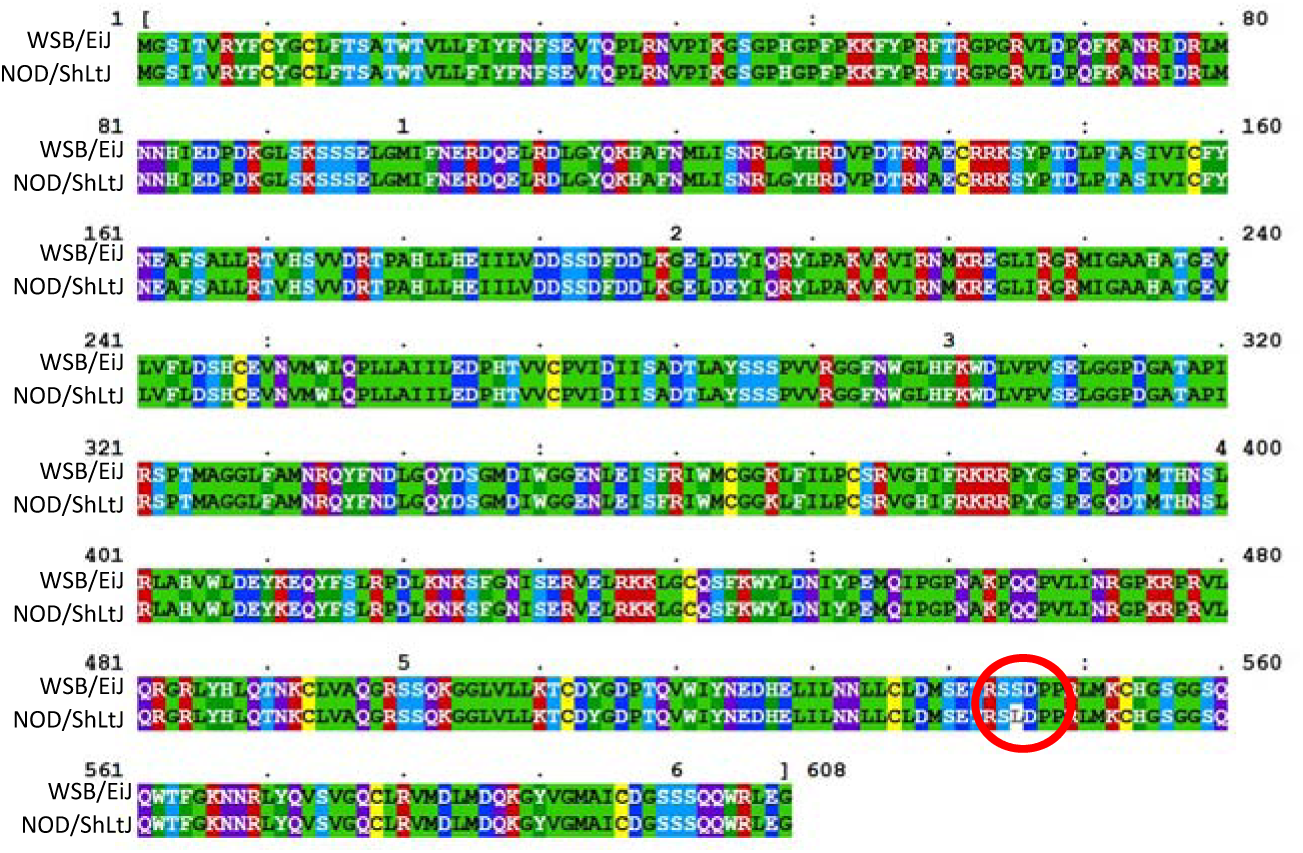
Protein sequence alignment between WSB/EiJ (WSB) and NOD/ShiLtJ (NOD) mice isoforms of GALNT11. The translated GALTNT11 protein sequence encoded by QTL driving alleles WSB and NOD demonstrate sequence homology except for S545L (*red circle*). Yellow=C (capable of disulfide bonding), Green=A, I, L, M, G, P, V (Hydrophobic) Dark Blue= D, E (Negative Charge), Dark Green =H, W, Y, F (Aromatic/Hydrophobic) Magenta= R,K (Positive Charge) light blue/purple=T, S, Q, N (Polar)

**Figure 7.**
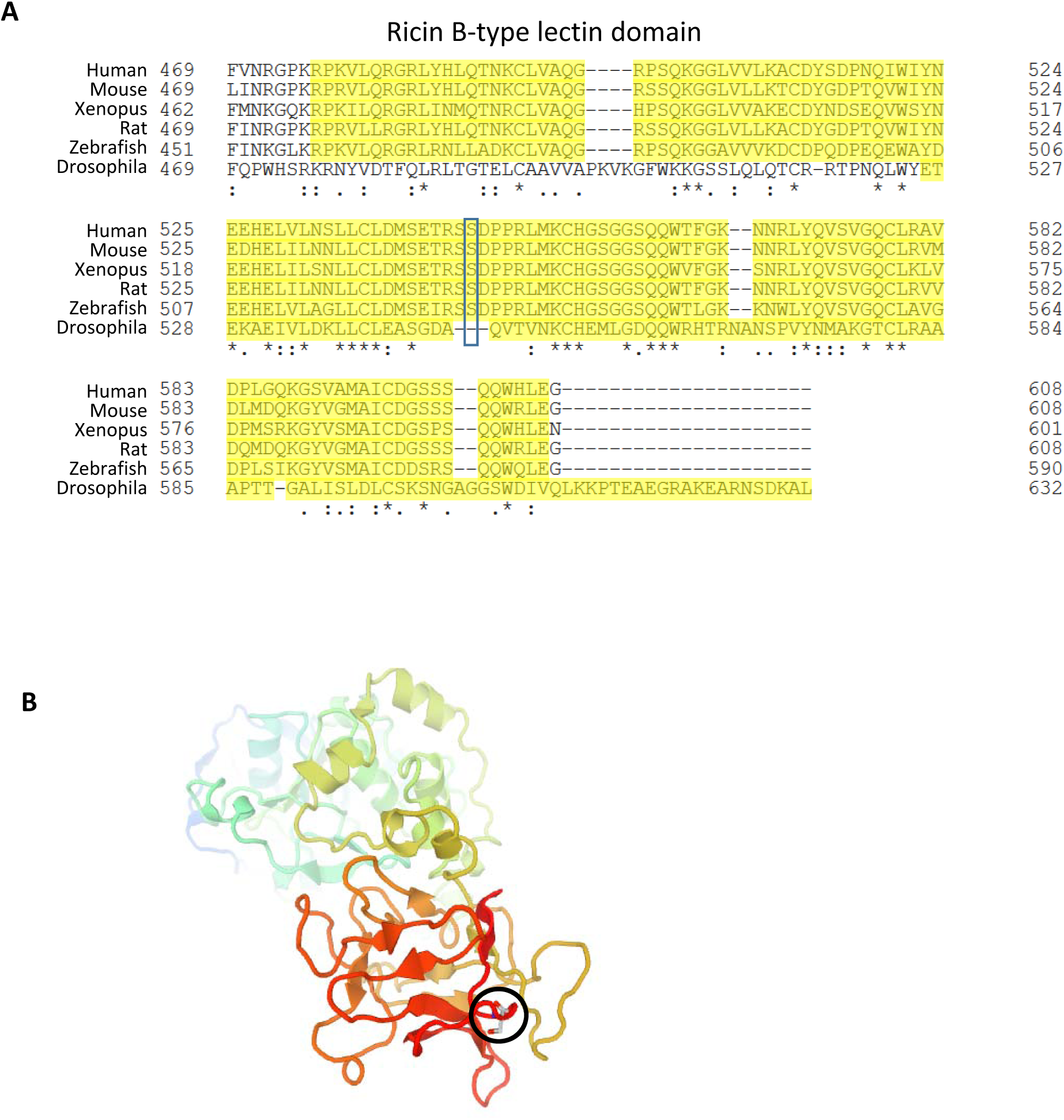
S545L occurs in the Ricin B lectin domain of GALNT11. (**A**) Multi-species alignment of the Ricin B lectin domain of GALNT11 showing that the S545L mutation occurs in a domain conserved across multiple vertebrates, including humans, mice, rats, *Xenopus*, and zebrafish. (**B**) Protein domain map of human GALNT11 showing the localization of the conserved S545L in the Ricin B lectin domain (*black circle*). The 3D structure rendered showing secondary structure as a cartoon type with coloring as a rainbow from N- to C-terminus.

### Integrative functional genomic analysis

*GALTN11* is a member of a family of an N-acetylgalactosaminyltransferases, which function in O-linked glycosylation. The non-synonymous serine-to-leucine SNP changes a conserved residue within the Ricin B lectin domain of GALNT11 and would be predicted to impede O-linked glycosylation. GALTN11 has been shown to uniquely glycosylate at least 313 glycoproteins in HEK cells^33^. Using the Jaccard similarity tool in the GeneWeaver^34^ system for integrative functional genomics, we determined that of the 313 human glycopeptides, 287 mouse orthologs are expressed in the mouse pre-Bötzinger complex^32^ (Table S2). Four of those genes *Hs6st2*^*35*^,*Fn1*^*36*^, *Lrp1*^*35*^, and *Sdc4*^*37*^ were identified by the Comparative Toxicogenomic Database^38^ as morphine-associated genes, and one additional pre-Bötzinger-expressed glycoprotein, *Cacna2d1*, has been shown to have differential brain expression in mice following morphine treatment^39^. Therefore, variants in *Galnt11* have a plausible mechanism of action through regulation of morphine-associated peptides in the pre-Bötzinger complex.

## Discussion

In this study, we found heritable strain differences in the quantitative metrics of respiratory response to morphine, including respiratory depression, recovery time and survival time, using an advanced, high-throughput, behavioral phenotyping protocol. We further identified genomic loci involved in morphine-induced respiratory depression using an unbiased genetic approach. Mapping these traits in the DO mice and evaluation of sequence variants and protein structure, followed by integrative functional genomic analysis in GeneWeaver, has allowed us to implicate *Galnt11* as a candidate gene for respiratory depression in response to morphine.

We identified specific inbred strains of mice that were more sensitive to morphine than other inbred strains of mice. The effect of sex was bidirectional in the inbred CC/DO founder strain population where some males and females differed from each other in either direction depending on strain. The traits of respiratory depression, recovery time and survival time were all shown to have a high degree of heritability and were largely independent traits. In determining our probe dose for the outbred population, we observed that the LD_50_ for morphine differed by four-fold between these eight parental strains harboring 45 million SNPs, or an equivalent genetic variation as found in the human population. Interestingly, mice such as AJ that had the lowest LD_50_ for both males and females, did not demonstate the highest degree of respiratory depression, suggesting that factors other than respiratory depression may play a role in opioid overdose.

To date, there have been no human GWAS or linkage studies for OIRD. Several GWAS studies have evaluated opioid use disorder (OUD) and/or opioid dependence, and have identified a variety of loci including SNPs linked to OPRM1, KCNG2, CNIH3, LOC647946, LOC101927293, CREB1, PIK3C3, and RGMA^40-45^. Additional human linkage studies have identified sex-specific traits^46^, epigenetic biomarkers^47^ or copy number variations^48^ for risk of OUD. Other studies have sought to separate opioid use from opioid dependence, and have thus far identified SNPs associated with SDCCAG8, SLC30A9, and BEND4^49^. Only one study has looked at human opioid overdose risk, specifically by scoring overdose status and determining the number of times that medical treatment was needed in European American populations^50^. In this study, SNPs near MCOLN1, PNPLA6 and DDX18 were identified as overdose risk alleles. Human genes have thus been mapped to opioid use, opioid dependence and opioid overdose susceptibility but human studies are not able to assess opioid-induced respiratory depression, specifically the LD_50_ of an opioid.

Animal studies have allowed us the opportunity to assess the LD_50_ of a drug in a variety of genetic backgrounds and then map those sources of variation. These types of controlled exposure experiments cannot be conducted in humans for which exquisite control of environment is not feasible and prior exposure history is unknown. Our genetic approach of QTL mapping in the DO mouse population has allowed us to identify a genomic region containing no genes previously known to function in opioid pharmacodynamics or pharmacokinetic processes, or implicated in OUD. The genetically diverse structure of this population allows for the identification of narrow genomic intervals often with very few candidate genes. This approach of using advanced mouse populations together with integrative functional genomics has been useful for the prioritization of candidate genes in a variety of different disciplines^51-53^

The identification of *Galnt11* as functioning within the morphine respiratory response reveals a potential new target for therapeutic development. GALNT11 is an N-acetylgalactosaminyltransferase that initiates O-linked glycosylation whereby an N-acetyl-D-galactosamine residue is transferred to a serine or threonine residue on the target protein. The lectin domain of GALNT11 is the portion that functions to recognize partially glycosylated substrates and direct the glycosylation at nearby sites. This type of post-translational modification controls many phamacokinetic and pharmacodynamic processes as well as the regulation of delta opioid receptor (OPRD1) membrane insertions as O-linked glycosylation is required for proper export of OPRD1 from the ER^54^. O-linked glycosylation is also required for opioid binding peptides, increasing their ability to cross the blood brain barrier^55^. The integrative functional analysis in GeneWeaver identified *Hs6st2*^*35*^,*Fn1*^*36*^, *Lrp1*^*35*^, and *Sdc4*^*37*^ as glycosylation targets of *Galnt11*. All are expressed in the pre-Bötzinger complex and are known to be responsive to morphine further supporting the concept that *Galnt11* is involved in morphine-related responses. Both *Hs6st2* and *Lrp1* were identified as increasing and decreasing, respectively, in responsive to both morphine and stress in C57BL/6J mice^35^. *Fn1* was upregulated six hours post morphine but downregulated four days later^36^, and *Sdc4*, a known-mu opioid receptor-dependent gene was also upregulated by morphine^37^. The drug-regulated expression of these known glycosylation targets of GALNT11 in relevant tissues further supports the functional relevance of *Galnt11* to OIRD.

Our findings demonstrate the initial mapping of a locus involved in OIRD in mice, for which the likely candidates do not act via the opioid receptor, thereby providing a potential new target for remedial measures. Although it is through mouse genetic variation that we identified this gene, it should be noted that this gene or its glycosylation targets need not vary in humans to be a viable target mechanism for therapeutic discovery and development. Characterization of the role of *Galnt11* and its variants along with other viable candidates will resolve the mechanism further, and continued mapping studies in larger populations will enable detection of additional loci for various aspects of the opioid induced respiratory response. These findings suggest that phenotypic and genetic variation in the laboratory mouse provides a useful discovery tool for identification of previously unknown biological mechanisms of OIRD.

## Methods

### Mice

Male (n=6) and female (n=6) mice from eight inbred strains including the founders that were used for the DO and CC population [C57BL/6J, 129S1/SvImJ, A/J, NOD/ShiLtJ, NZO/HlLtJ, /CAST/EiJ, PWK/PhJ, WSB/EiJ] were tested at each dose of morphine. Male and female DO mice (n =300 including 150 of each sex; J:DO, JAX stock number 009376) from generation 28 of outcrossing were used. All mice were acquired from The Jackson Laboratory (JAX) and were housed in duplex polycarbonate cages and maintained in a climate-controlled room under a standard 12:12 light-dark cycle (lights on at 0600 h). Bedding was changed weekly and mice had free access to acidified water throughout the study. Mice were provided free access to food (NIH31 5K52 chow, LabDiet/PMI Nutrition, St. Louis, MO). A Nestlet and Shepherd Shack were provided in each cage for enrichment. Mice were housed in same sex groups of three to five mice per cage. All procedures and protocols were approved by JAX Animal Care and Use Committee and were conducted in compliance with the NIH Guidelines for the Care and Use of Laboratory Animals.

### Morphine

Morphine sulfate pentahydrate (NIDA Drug Supply) was prepared at varying concentrations in sterile saline to deliver doses (200-1200 mg/kg s.c.) in a manner not to exceed 0.2 ml/10g body weight. Starting with a dose of 436 mg/kg for all strains and depending upon the result of that dose, the next dose was either increased (536 mg/kg) or decreased (336 mg/kg) for the next cohort, such that doses flanking both sides of the 50% survival point (LD_50_) were tested. This was repeated with increasing and decreasing doses as necessary depending upon the results of the previous dose. Not all strains received all doses but each strain received at least three doses such that two flanked (one above, one below) the LD_50_. Using this approach and testing 3-4 doses per strain, the LD_50_ for the eight strains and both sexes was determined and ranged from 212 mg/kg - 882 mg/kg.

### Piezoelectric sleep monitoring to determine respiratory depression, recovery time and survival time

Mice were placed individually into the 7 x 7 inch piezoelectric grid and chamber system for 24 hours to equilibrate to the apparatus and collect baseline activity data^26,56^. Individual testing is necessary due to the known enhanced lethality of cage mates during morphine exposure, which has been shown to affect survival^57^. The mice had access to food and water *ad libitum* while in the chamber. The room was maintained on a 12:12-h light:dark cycle. To control for known circadian effects^58^ mice were placed in the chambers between 9-12 am on Day 1 and were injected with morphine 24 hours later. They remained in their chambers undisturbed until 24 hours after injection. Whenever possible, complete balanced cohorts of the eight strains and both sexes were run during each of nine replicates of the experiment. The data acquisition computer, food and water were checked daily; otherwise, the mice remained undisturbed. Breath rates were estimated from 4-second intervals in which animal activity dropped low (i.e. during sleep and brief rest periods and pauses during wake), and averaged over 24-minute overlapping intervals to provide an average respiratory rate every 12 minutes. The respiratory rate baseline consisted of the average respiratory rate over the first 24 hours, which included both sleep and wake periods. Respiratory rate was then measured in the same way after injection of opioid. These measures were then used to determine thresholds for obtaining the recovery time (respiratory rate returns to baseline, see Fig. 1) or survival time (animal stops moving and breathing, never returning to threshold, see Fig. 1). The 300 DO mice were tested in random cohorts of 36 using the PiezoSleep system with a single 486 mg/kg dose of morphine. This dose was determined as the average LD_50_ dose across the eight strains and two sexes, 16 samples.

### Calculating respiratory depression, recovery time and survival time

To test for difference in the respiratory depression, recovery time and survival time across the strains and sexes a linear model was fit, the full model was:

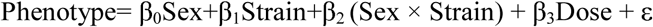

where Phenotype was respiratory depression, recovery time or survival time and where ε is random error. The β-parameters were estimated by ordinary least squares, and the type III sum of squares wa considered ε in the ANOVA model. In all cases, the full model was fit and reduced by dropping non-significant interactions followed by main effects.

### Calculating broad sense heritability (H2)

As a measure of broad sense heritability in the founder strains, the ICC was determined using ICCest from the ICC 2.3.4^59^ package in R 4.0.0.

### Morphine LD_50_ data analysis

The LD_50_ was calculated using the drc 3.0-1 library^60^ in R using the total tested and the observed dead at each dose as a binary or binomial response. A logistic regression model was fit, and a goodness of fit test (based upon Bates and Watts^61^) performed. In addition, a regression model assuming equal LD_50_ across strains was compared by chi square to an LD_50_ assumed different across strains. An adjusted and unadjusted 95% credible interval was also calculated. To graph the data a non-linear 2-parameter model was fit in JMP 14.2.0 (RRID:SCR_014242) with the formula:

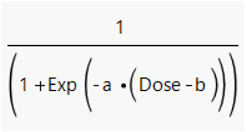

where a=growth rate and b=inflection point.

### Genotyping

Tail samples were collected at the conclusion of the experiment and all mice were genotyped using the Giga-MUGA genotyping array (NeoGene). Data were deposited in the DO Database (DODB) RRID:SCR_018180). Genotypes were imputed to a 69K grid to allow for equal representation across the genome.

### QTL Mapping

Genotype probabilities were calculated according to the founder genotypes and then converted to allele probabilities. We then interpolated allele probabilities into a grid of 69,000 evenly-spaced genetic intervals^62^. We performed genome scans using R/qtl2 RRID:SCR_018181^63^. Sex and date of test were included as additive covariates. The model includes the random effect of kinship among the DO animals computed using the LOCO method^64^. The significance thresholds were determined for each trait by permutation mapping^65^. The confidence interval around the peak makers was determined using Bayesian support intervals. To determine the percent of variation accounted for by the QTL the mice were classified at the variant into one of eight states based on genotype probabilities of mice at that locus. Following this, a one-way ANOVA was fit to ascertain the strain variation relative to total variation towards estimating heritable variation at that locus. SNP association mapping was also performed using R/qtl2 to test the association of individual SNPs alleles in the region of the locus with the phenotype. Briefly, SNP data were obtained from the SANGER and MGI databases^66,67^ for the interval. Using the genotype probabilities and the founder SNP genotypes to infer the SNP genotypes of the DO mice. At each SNP location the eight allele state probabilities are collapsed to two state SNP probabilities are collapsed to The Cox proportional hazards regression was performed by coxph function in the survival (3.1-12) R package. Based on the output of the log (base e) likelihood for the null model and for the alternative model (with covariates and genotype probabilities), we took the difference of both log likelihoods and then divided by ln(10) to convert the results into the LOD scale. Full QTL mapping scripts are available https://thejacksonlaboratory.github.io/DO_Opioid/index.html

### Candidate gene analysis

In order to assess the plausibility of genes in the QTL interval we identified all SNPs within the additive SNP model segregating between the high and low alleles of the DO founder strains. Next we identified those that were within protein coding region that were most deleterious. Differential coding sequence non-synonymous amino acid substitution SNPs (Cn) that differed between the high and low allele groups were identified. GeneWeaver’s database (RRID:SCR_003009) was searched to identify the overlap among tissue-specific expression profiles from Allen Brain Atlas as well as datasets derived from Entrez GEO profiles (RRID:SCR_004584) for pre-Bötzinger neurons^32^ (GS 273275), diaphragm (GS273269)^68^ and lung^69^ with the QTL positional candidates,. The gene sets were overlapped using the Jaccard similarity and GeneSet graph tools^34^.

In order to determine if the Cn SNPS were in areas of evolutionary conservation we aligned the sequence of several species. Representative sequences for each species *Drosophila* (*Q8MVS5), Xenopus (Q6DJR8), Danio (*Q08CC3), *Rattus (Q6P6V1), Mus (Q921L8), Homo* (*Q8NCW6*) were acquired from Uniprot (RRID:SCR_002380) and aligned. The clustalo program was used with default parameters^70^. The transition matrix is Gonnet, gap opening penalty of six bits, gap extension of one bit. Clustal-Omega uses the HHalign algorithm and its default settings as its core alignment engine^71^. To determine where the three-dimensional effects of the amino acid change would be we obtained the 3D crystal structure (1XHB)^72^ from the Research Collaboratory for Structural Bioinformatics Protein Data Bank (RRID:SCR_012820) and visualized it with Jmol (RRID:SCR_003796)^73^.

### Integrative functional genomics

In order to assess the functional sufficiency of *Galnt11* as a candidate gene the literature was searched to identify genome-wide studies characterizing glycosylation targets of GALTN11, one study was identified^33^ and these genes were added to the GeneWeaver Database (GS356053). Using the Jaccard similarity tool, we overlapped the glycosylation targets with the genes expressed in the mouse pre-Bötzinger complex. We next overlapped these gene with genes identified by the Comparative Toxicogenomic Database (RRID:SCR_006530) as morphine-associated genes.

## Supporting information

Supplemental Tabl 1

## Figure Legends

**Figure S1.**
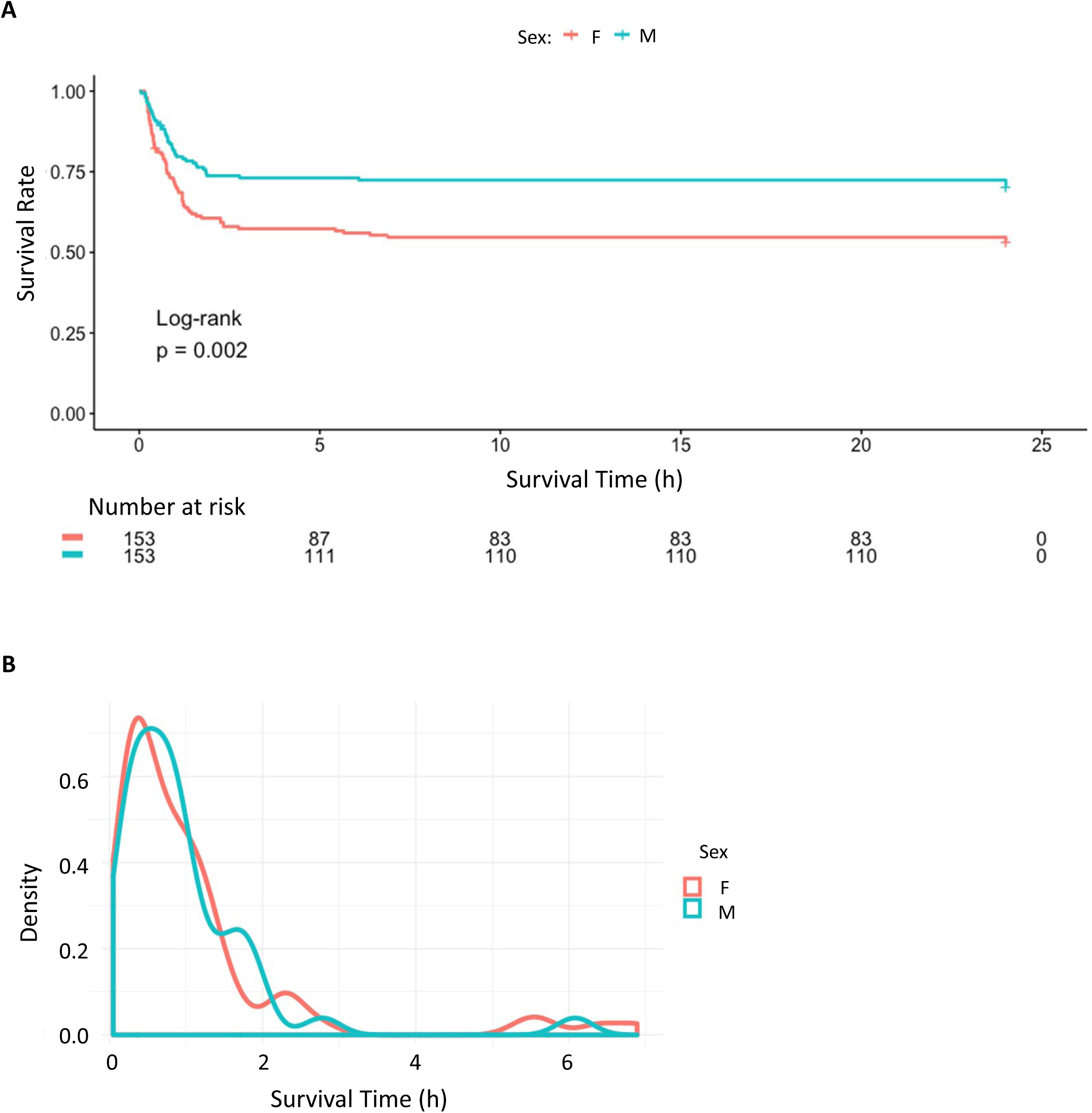
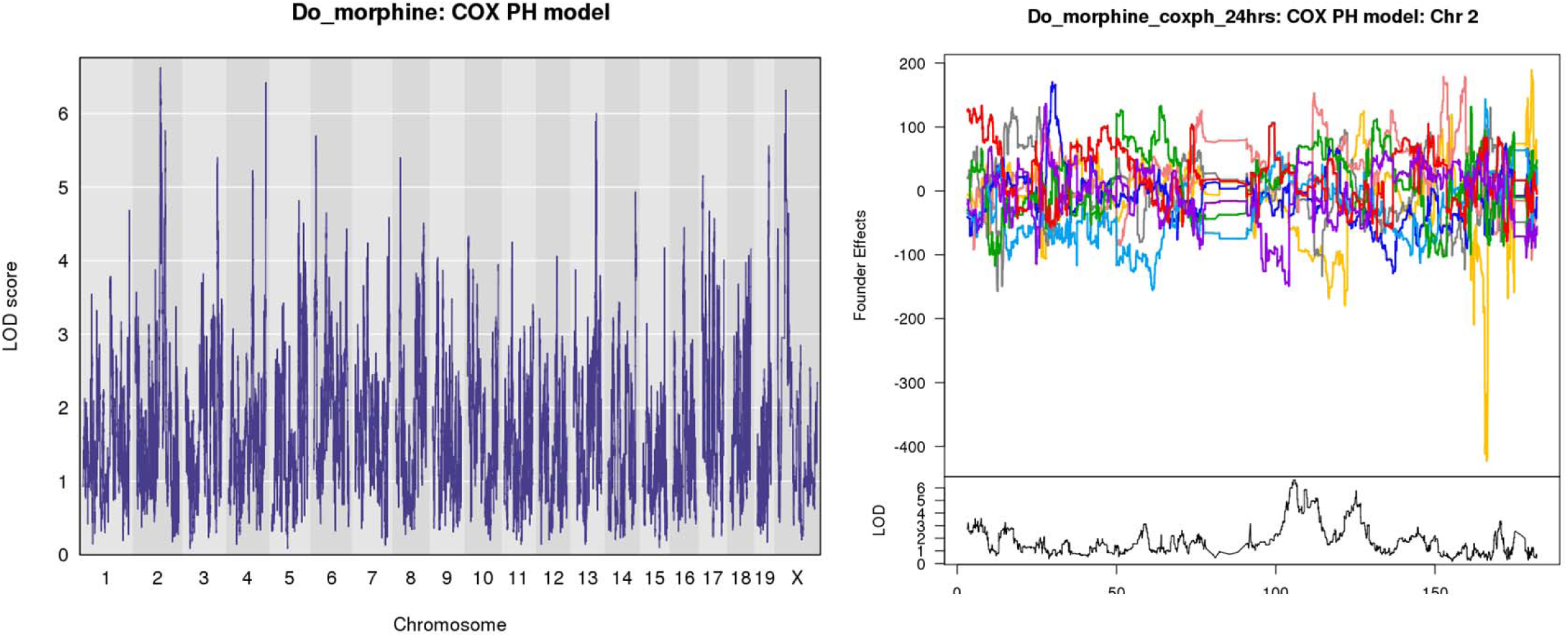
Mapping of the DO survival data using a Cox Proportional-Hazards Model. (**A**) Survival time of male (M, blue) and female (F, red) DO mice as represented by a Cox Proportional-Hazards Model. (**B**) The difference of these log likelihoods was taken and then divided by ln (10) to convert the result to the LOD scale. (**B**) Cox Proportional Hazards (COXPH) QTL mapping model which includes the genotype probabilities. (**D**) Allele effect plot of suggestive chromosome 2 locus.

**Table S1**. The 1,885 SNPs that differ between NOD and WSB within the Chromosome 5 QTL interval.

**Table S2**. The 287 genes that are known targets for GALNT11 and expressed in the pre-Botzinger complex

## Acknowledgements

Funding R01 DA037927, R01 AA018776 and P50 DA039841to EJC, R01 DA048890 to JAB, and the NIDA Drug Supply Program. Special thanks to Jennifer Ryan and the Center for Biometric analysis at JAX for use of the PiezoSleep system and to Stephen Krasinski for extensive edits of the manuscript.

